# FINCA disease mouse model exhibits altered behaviour and immune response

**DOI:** 10.1101/2024.06.14.599017

**Authors:** Anniina E. Hiltunen, Salla M. Kangas, Aishwarya Gondane, Henna Koivisto, Kari Salokas, Anne Heikkinen, Miia H. Salo, Tapio Röning, Antti Tallgren, Virpi Glumoff, Maria C. Denis, Niki Karagianni, Johanna Myllyharju, Markku Varjosalo, Heikki Tanila, Harri M. Itkonen, Mika Rämet, Johanna Uusimaa, Reetta Hinttala

**Affiliations:** Medical Research Center Oulu and Research Unit of Clinical Medicine, Faculty of Medicine, University of Oulu and Oulu University Hospital, Oulu, Finland; Biocenter Oulu, University of Oulu, Oulu, Finland; Department of Biochemistry and Developmental Biology, Faculty of Medicine, University of Helsinki, Helsinki, Finland; A.I. Virtanen Institute for Molecular Sciences, University of Eastern Finland, Kuopio, Finland; Institute of Biotechnology, Helsinki Institute of Life Science (HiLIFE), University of Helsinki, Helsinki, Finland; ECM-Hypoxia Research Unit, Faculty of Biochemistry and Molecular Medicine, University of Oulu, Oulu, Finland; Medical Research Laboratory Unit, Faculty of Medicine, University of Oulu, Oulu, Finland; Biomedcode Hellas SA, Vari, Athens, Greece; Faculty of Medicine and Health Technology, Tampere University, Tampere, Finland; Department of Children and Adolescents, Division of Paediatric Neurology, Oulu University Hospital, Oulu, Finland

**Keywords:** FINCA disease, NHLRC2, NHL repeat containing 2, neurodevelopmental disorder, immune response, mouse model

## Abstract

Fibrosis, neurodegeneration and cerebral angiomatosis (FINCA) is a childhood-onset multi-organ neurodevelopmental disorder associated with multi-organ manifestations and recurrent infections. The disease is caused by variants in *NHLRC2* initiating a cascade of unknown pathological events. Previously, we have demonstrated that despite the significant decrease at the molecular level, the compound heterozygosity of knock out and p.Asp148Tyr alleles in NHLRC2 does not lead to a severe phenotype in mice. Here, we analysed the behavioural and immunological phenotype of the FINCA mice and studied the molecular pathways affected by p.Asp148Tyr in NHLRC2 using mouse and human-derived cell culture models. The FINCA mice displayed a mild hyperactivity and deficient early immune response when challenged with LPS leading to altered cytokine responses, including IFNγ, IL-12, and TNFα. By comparing gene expression and putative interaction partners affected by p.Asp148Tyr, we identified Rho GTPase signalling as the common pathway. Altogether, these results establish a multi-dimensional impact of the p.Asp148Tyr variant in NHLRC2. Knowledge of the molecular pathways affected by NHLRC2 and the natural course of FINCA disease progression are instrumental for the development of effective therapeutics.

**Summary statement:** FINCA is a paediatric neurodevelopmental and multi-organ disorder caused by variants in *NHLRC2*. Here, mild hyperactivity in connection with altered early immune response is described in the FINCA mouse model.

## Introduction

Fibrosis, neurodegeneration, and cerebral angiomatosis (FINCA) (OMIM 618278) is a rare infantile-onset disease caused by variants in the NHL-repeat-containing 2 (*NHLRC2*) gene. The disease is characterized by neurodevelopmental disorder, intellectual disability, progressive neurodegeneration, tissue fibrosis, infection susceptibility/immunodeficiency and chronic haemolytic anaemia (Tallgren et al., 2023; Uusimaa et al., 2018). Since our initial characterisation, 33 patients from various ethnic backgrounds have been reported with different combinations of recessive pathogenic variants in NHLRC2 broadening the phenotypic spectrum of the disease (Badura-Stronka et al., 2022; Brodsky et al., 2020; Li et al., 2022; Rapp et al., 2021; Sczakiel et al., 2023; Tallgren et al., 2023). Thus far, all reported FINCA patients presented with developmental delay and variable disease progression with or without pulmonary involvement. In some patients, recurrent infections alleviated later in life, and severe neurodevelopmental disorder evolved as the main clinical feature. To date, our understanding of the pathomechanism of the disease is limited; thus, treatment relies on symptom-alleviating options.

Previous studies have implicated NHLRC2 in vesicle transport and cytoskeleton organisation in human dermal fibroblasts (Paakkola et al., 2018) as well as oxidative stress regulation in colon cancer cells (Nishi et al., 2017). Moreover, in human macrophages, NHLRC2 has been identified as a regulator of phagocytosis, potentially influencing actin dynamics through RhoA-Rac1 signalling (Haney et al., 2018; Yeung et al., 2019). Structurally, the NHLRC2 protein comprises three domains: an N-terminal thioredoxin-like (Trx-like) domain, a six-bladed NHL-repeat-containing β-propeller domain, and a C-terminal β-stranded domain. Pathogenic variants have been identified across all domains, with the most prevalent FINCA disease-causing variant, p.Asp148Tyr, residing in the Trx-like domain. Although the structure of NHLRC2 suggests an enzymatic role for the protein, its detailed molecular function remains unknown (Biterova et al., 2018).

To study the pathophysiology of FINCA disease, we previously generated a FINCA mouse model harbouring the NHLRC2 variant p.Asp148Tyr in one allele and *Nhlrc2* knockout in the other allele (henceforth referred to as FINCA mouse), similar to the genotype of first identified Finnish patients. We confirmed the disturbance of the vesicular transport pathway and actin dynamics-associated proteins in neuronal precursor cells (NPCs) due to the p.Asp148Tyr variant in these mice (Hiltunen et al., 2020). In addition, we discovered an increased amount of isoform two of the heterogeneous nuclear ribonucleoproteins C1/C2 (*Hnrnpc*, hnRNPC2) in NPCs as well as the hippocampus of the FINCA mouse model, indicating a possible involvement of RNA homeostasis in the disease course (Hiltunen et al., 2020). Despite observing a significant decrease in NHLRC2 in all tissues and alterations in neuronal proteins, these mice do not exhibit the severe symptoms characteristic of FINCA disease (Hiltunen et al., 2020). We hypothesised that the absence of immunological challenges in these mice may protect them from FINCA disease progression, given that patients often experience worsening symptoms post infection.

Here, we explore potential changes in RNA homeostasis by examining transcriptional and splicing changes in both the FINCA variant and *Nhlrc2* knockout (KO) homozygous mouse embryonic stem cells (mESCs). We perform detailed secondary phenotyping of our FINCA mouse model, focusing on their behavioural phenotypes and immunological responses. Additionally, we translate our findings from mice to human cells through *in vitro* studies. The objective is to explore the molecular pathways underlying FINCA and to enable the development of interventions that ameliorate the disease course in the future.

## Results

### Transcriptional changes in *Nhlrc2* KO and p.Asp148Tyr variant mESCs compared to wild type cells

To date, mESCs are the only known cell type to tolerate the complete loss of NHLRC2 (Hiltunen et al., 2022). We utilised this feature to examine transcriptional changes in *Nhlrc2* KO mESCs to gain more information about the biological role of NHLRC2. Furthermore, mESCs homozygous for p.Asp148Tyr in NHLRC2 were established and studied to better understand the consequences of the FINCA disease-causing variant. RNA sequencing was performed on four wild-type (WT), four homozygous *Nhlrc2* KO and four homozygous p.Asp148Tyr variant (FINCA variant) mESC lines. In the KO mESCs, the transcription of the *NHLRC2* gene is halted by a trapping cassette between exons 4 and 5 (Hiltunen et al., 2022; Skarnes et al., 2011). The FINCA variant mESCs express full-length endogenous *Nhlrc2* mRNA modified to contain the c.442G > T variant, leading to p.Asp148Tyr as well as a silent c.408C>A for genotyping purposes (Hiltunen et al., 2020).

Our analysis revealed 74 differentially expressed genes (DEGs) with an adjusted p-value (padj) of less than 0.05 in KO cells and 3712 DEGs in FINCA variant cells compared to WT cells (Fig. 1A, Suppl. Data 1). The FINCA variant cells presented statistically significant changes in 79.7% of the 74 DEGs detected in KO cells (Fig. 1B). All but two of the common DEGs (*Tlcd4, S1pr1*) were changed to the same direction. To identify the major biological processes affected, we performed KEGG and REACTOME pathway analysis of the FINCA variant DEGs. This analysis indicated signalling pathways of metabolism, lysosomes, cell proliferation, cell adhesion and interaction with extracellular matrix, collagen metabolism, neurodevelopment and axonal growth, Ras and Rho GTPase signalling, cancer, and epigenetic regulation to be affected (Table 1). In KO DEGs, gene ontology (GO) term enrichment analysis indicated an association with cell differentiation and developmental processes (Table 1) which is in agreement with our earlier data showing that NHLRC2 is essential for early embryogenesis (Hiltunen et al., 2022).

**Figure 1.**
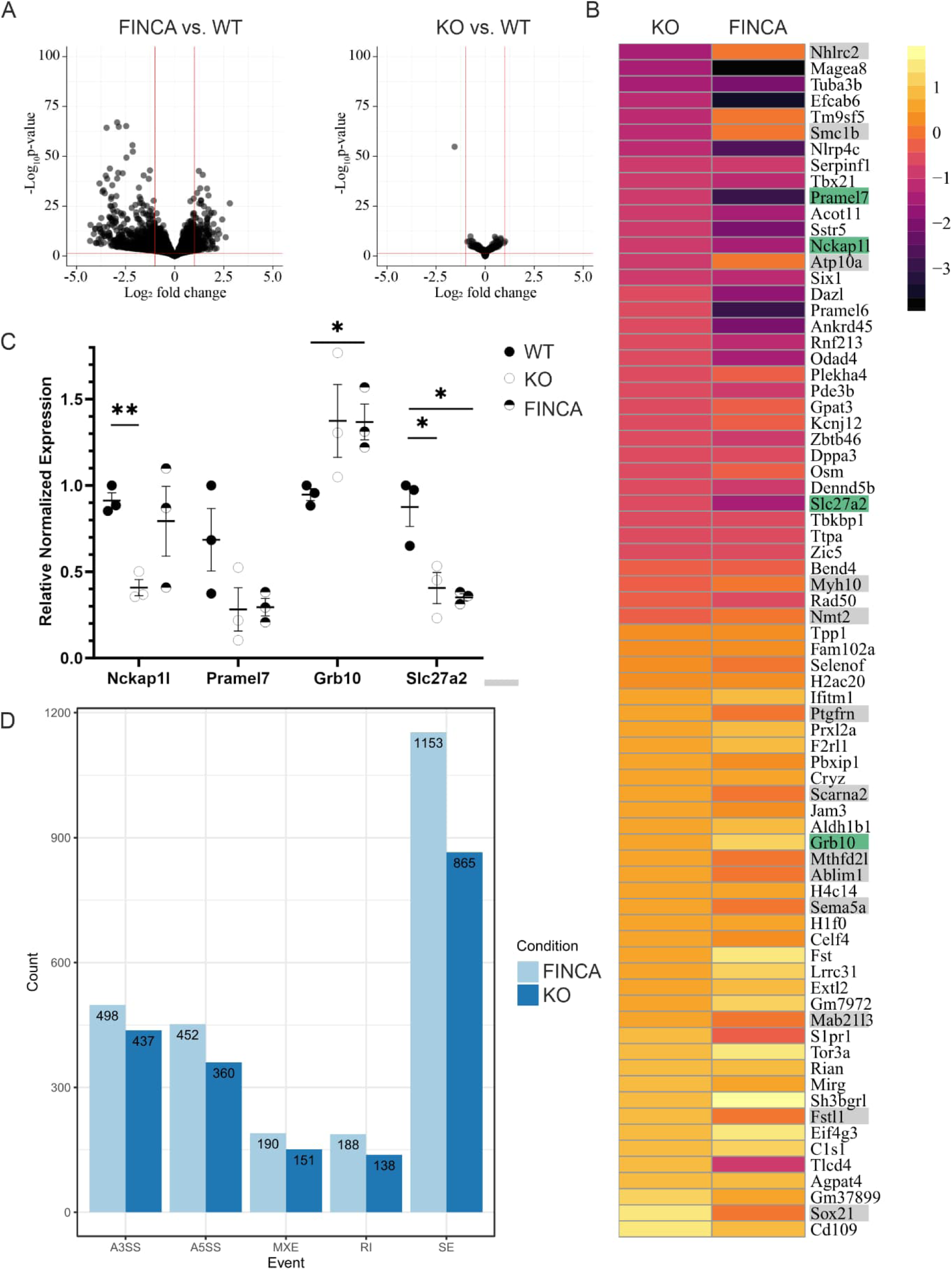
RNA sequencing of homozygous FINCA variant and Nhlrc2 KO in mouse embryonic stem cells. A. Volcano blot of the FINCA variant and Nhlrc2 KO DEGs show more profound overall changes in the FINCA variant vs. WT comparison compared to the Nhrlc2 KO vs. WT comparison. Significant genes have padj-value < 0.05. B. Gene expression heatmap of the DEGs in Nhlrc2 KO and FINCA variant mESCs compared to WT cells (group means). Grey genes showed no statistically significant difference between the FINCA variant and WT mESCs. Green genes were selected for validation. C. qPCR verification of Nckap1l, Pramel 7, Grb10 and Slc27a2. Statistical analyses were conducted using an unpaired Student’s t-test, * p < .05, **p < 0.01. Scatter plots show the individual data points, group means, and standard error of means (SEM). Expression was relative to one of the WT samples. Glyceraldehyde 3-phosphate dehydrogenase (Gapdh) and actin beta (Actb) were used as reference genes. D. Count of different alternative splicing events in FINCA and KO mES cells compared to WT cells (p-value < 0.05). Main types of alternative splicing events: A3SS: alternative 3′ splice sites, A5SS: alternative 5′ splice sites, MXE: mutually-exclusive exons, RI: retained intron and SE: exon skipping.

**Table 1.**
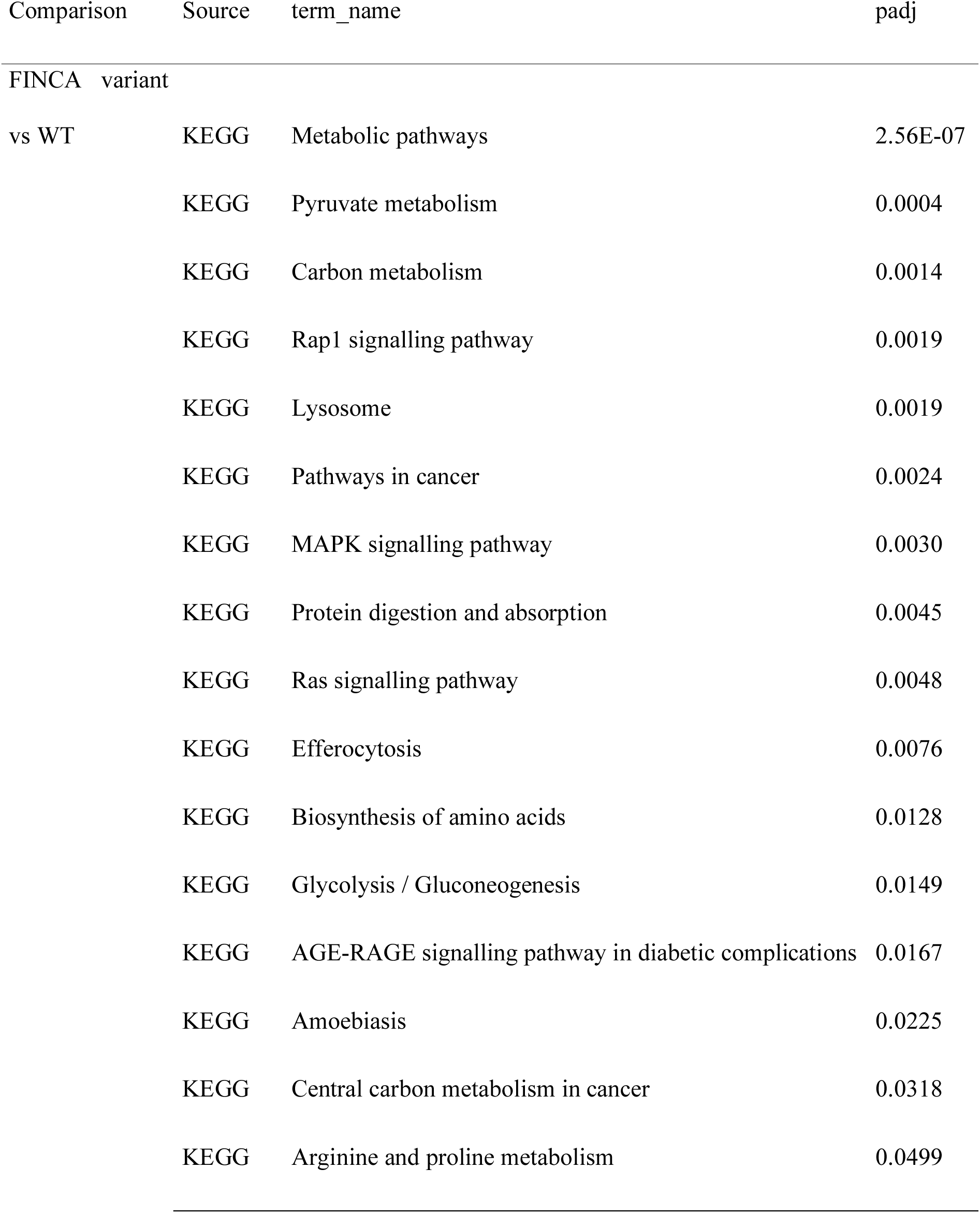
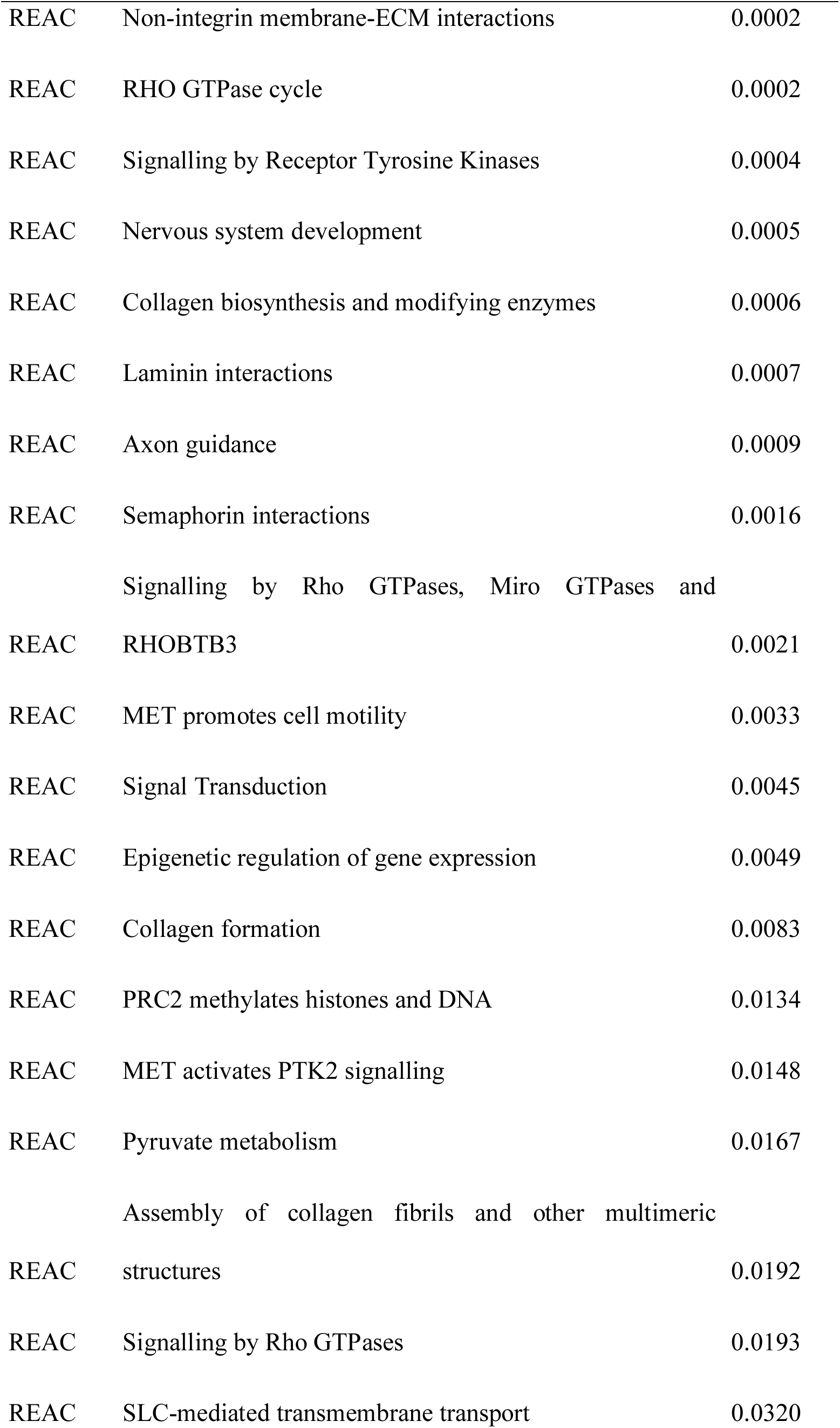
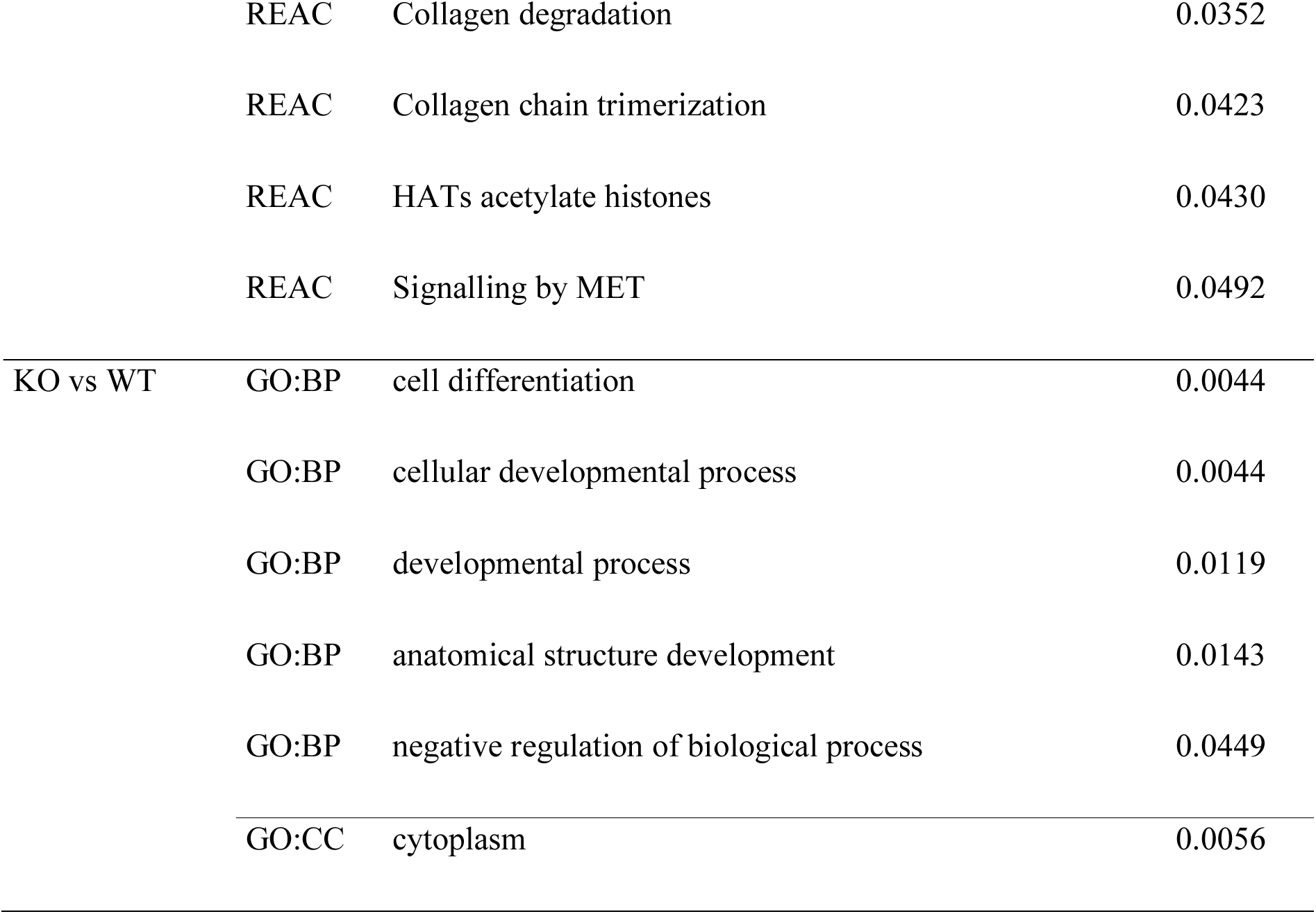
Pathway analysis of the FINCA variant vs. WT and GO term enrichment analysis of KO vs. WT DEGs identified in mESCs. The organism used in the analysis was Mus musculus.

| Comparison | Source | term_name | padj |
| --- | --- | --- | --- |
| FINCA variant |  |  |  |
| vs WT | KEGG | Metabolic pathways | 2.56E-07 |
|  | KEGG | Pyruvate metabolism | 0.0004 |
|  | KEGG | Carbon metabolism | 0.0014 |
|  | KEGG | Rap1 signalling pathway | 0.0019 |
|  | KEGG | Lysosome | 0.0019 |
|  | KEGG | Pathways in cancer | 0.0024 |
|  | KEGG | MAPK signalling pathway | 0.0030 |
|  | KEGG | Protein digestion and absorption | 0.0045 |
|  | KEGG | Ras signalling pathway | 0.0048 |
|  | KEGG | Efferocytosis | 0.0076 |
|  | KEGG | Biosynthesis of amino acids | 0.0128 |
|  | KEGG | Glycolysis / Gluconeogenesis | 0.0149 |
|  | KEGG | AGE-RAGE signalling pathway in diabetic complications | 0.0167 |
|  | KEGG | Amoebiasis | 0.0225 |
|  | KEGG | Central carbon metabolism in cancer | 0.0318 |
|  | KEGG | Arginine and proline metabolism | 0.0499 |

|  |  |  |
| --- | --- | --- |
| REAC | Non-integrin membrane-ECM interactions | 0.0002 |
| REAC | RHO GTPase cycle | 0.0002 |
| REAC | Signalling by Receptor Tyrosine Kinases | 0.0004 |
| REAC | Nervous system development | 0.0005 |
| REAC | Collagen biosynthesis and modifying enzymes | 0.0006 |
| REAC | Laminin interactions | 0.0007 |
| REAC | Axon guidance | 0.0009 |
| REAC | Semaphorin interactions | 0.0016 |
|  | Signalling by Rho GTPases, Miro GTPases and |  |
| REAC | RHOBTB3 | 0.0021 |
| REAC | MET promotes cell motility | 0.0033 |
| REAC | Signal Transduction | 0.0045 |
| REAC | Epigenetic regulation of gene expression | 0.0049 |
| REAC | Collagen formation | 0.0083 |
| REAC | PRC2 methylates histones and DNA | 0.0134 |
| REAC | MET activates PTK2 signalling | 0.0148 |
| REAC | Pyruvate metabolism | 0.0167 |
|  | Assembly of collagen fibrils and other multimeric |  |
| REAC | structures | 0.0192 |
| REAC | Signalling by Rho GTPases | 0.0193 |
| REAC | SLC-mediated transmembrane transport | 0.0320 |
|  | REAC | Collagen degradation | 0.0352 |
|  | REAC | Collagen chain trimerization | 0.0423 |
|  | REAC | HATs acetylate histones | 0.0430 |
|  | REAC | Signalling by MET | 0.0492 |
| KO vs WT | GO:BP | cell differentiation | 0.0044 |
|  | GO:BP | cellular developmental process | 0.0044 |
|  | GO:BP | developmental process | 0.0119 |
|  | GO:BP | anatomical structure development | 0.0143 |
|  | GO:BP | negative regulation of biological process | 0.0449 |
|  | GO:CC | cytoplasm | 0.0056 |

We validated the RNA sequencing results by selected DEGs from a variety of cellular processes but common to both the FINCA variant and KO mESCs using quantitative PCR (qPCR) (Fig 1B, C). The selection was based on the degree of dysregulation and expression abundance in RNA sequencing analysis in both genotypes. Three new mESC samples were grown and prepared for the analysis from each genotype. Nck-associated protein 1-like (*Nckap1l* aka *Hem1*) was significantly decreased in KO cells compared to WT cells (0.91 vs. 0.41, p = 0.0015). We noted a modest but non-significant decrease in the expression of *Nckap1l* also in FINCA variant cells. PRAME-like 7 (*Pramel7*) was reduced, but not statistically significantly, in KO and FINCA variant cells compared to WT cells. Growth factor receptor-bound protein 10 (*Grb10*) expression was increased in FINCA variant cells compared to WT cells (0.946 vs. 1.368, p = 0.018) and although not statistically significant, a similar trend was seen in KO cells. Solute carrier family 27 (fatty acid transporter), member 2 (*Slc27a2*) was significantly decreased in both KO (0.875 vs. 0.406, p = 0.031) and FINCA variant (0.875 vs 0.352, p = 0.0102) cells compared to WT cells. Altogether, three out of four selected genes had a statistically significant change (p < 0.05) to the same direction in the mRNA expression level either in one (KO) or both (KO and FINCA variant) cell cultures, validating the results from RNA sequencing.

Previously, we have connected NHLRC2 dysfunction with the accumulation of RNA binding protein, hnRNP C2, in developing neurons and the hippocampus of adult female FINCA mice (Hiltunen et al., 2020). To further probe the possibly disturbed RNA homeostasis due to the homozygous FINCA variant in mESCs, we performed alternative splicing analysis to identify the five major alternative splicing categories: alternative 3′ and 5′ splice sites (A3’SS and A5’SS, respectively), mutually exclusive exons (MXE), detained introns (DI) and skipped exons (SE) using rMATS (Shen et al., 2014). This approach identified 2481 significant alternative splicing events in the FINCA variant cells and 1951 in the KO cells (Fig. 1D). However, the magnitude of the change was modest, even for the most significant events (Fig. S1). In summary, the alternative splicing analysis indicated that the mESCs cultured *in vitro* were able to adapt to both the homozygous p.Asp148Tyr variant and loss of NHLRC2. However, it is possible that splicing becomes important as cells differentiate or respond to stress — for example, when challenged with a pathogen.

### FINCA mice show ADHD-like phenotype

As neurodevelopmental delay is one of the main manifestations of FINCA disease (Badura-Stronka et al., 2022; Brodsky et al., 2020; Li et al., 2022; Rapp et al., 2021; Sczakiel et al., 2023; Tallgren et al., 2023; Uusimaa et al., 2018), we conducted a behavioural neurophenotyping of FINCA mice and their WT littermates at 2.5 months and again at 9 months of age. At 2.5 months, the FINCA mice showed increased ambulatory distance (1362 cm vs. 1677 cm, p = 0.015) and decreased mean rest time (124 s vs 94 s, p = 0.03) in the TruScan apparatus compared to the WT littermates (Fig 2A). In the novel object recognition task, the FINCA mice spent less overall time exploring the test objects (58 s vs. 43 s, p = 0.04) than the WT mice (Fig. 2B). No significant genotype differences were observed in nestbuilding, elevated plus maze, the marble burying task, grid hanging, social approach, rotarod, hot plate, novelty-suppressed feeding, the Morris swim task or forced swimming tests at 2.5 months of age. No differences in behavioural tests were recognised at 9 months of age, including ambulatory distance and rest time in the TruScan. Notably, the ambulatory distance of the WT mice was about the same (1326 cm) as at 2.5 moths, while the FINCA mice moved substantially less (1308 cm) than at a younger age. Collectively, these findings suggest that FINCA mice have mild hyperactivity during adolescence, which abates by full adult age, resembling attention deficit hyperactivity disorder (ADHD)-like behaviour.

**Figure 2.**
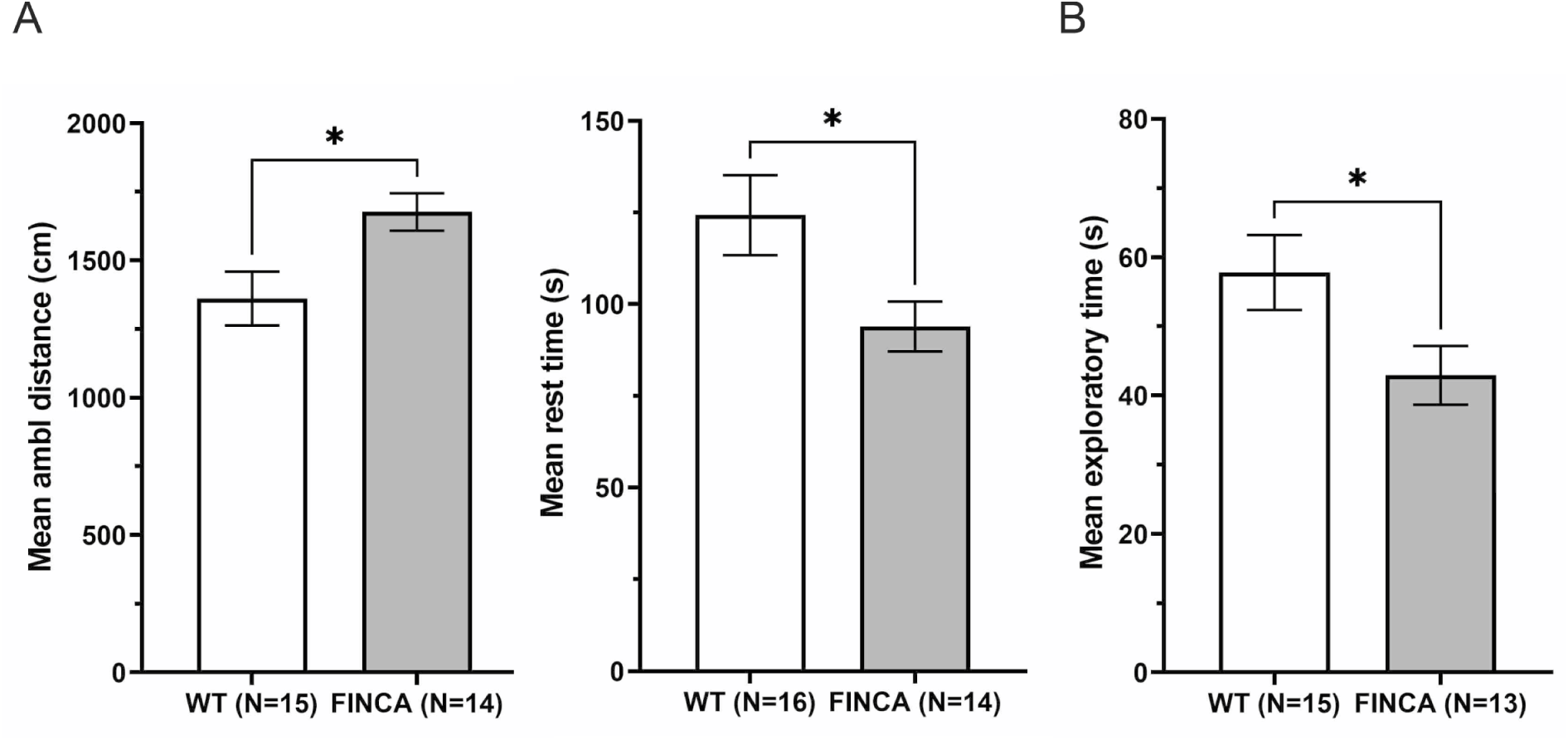
Behavioral tests of FINCA mice and WT littermates at 13 weeks of age. A. Spontaneous exploratory activity test revealed increased ambulatory distance and decreased mean rest time in the the FINCA mice compared to the WT mice. B. FINCA mice spent less time exploring a novel object than WT mice. An unpaired Student’s t-test was used for the statistical analysis. Group means and SEMs are shown. * p < 0.05.

### Early innate immune response is altered in FINCA mice

Another common manifestation of FINCA disease is recurrent infections (Badura-Stronka et al., 2022; Brodsky et al., 2020; Li et al., 2022; Rapp et al., 2021; Sczakiel et al., 2023; Tallgren et al., 2023; Uusimaa et al., 2018), suggesting that NHLRC2 deficiency affects the immune system. To evaluate the innate immune response of FINCA mice, we used lipopolysaccharide (LPS) induction and measured 13 common cytokines as a first screen from serum samples obtained 1.5 and 6 h post-induction from 8-week-old mice. The FINCA mice showed decreased serum interferon γ (IFNγ) (2.2 pg/ml vs. 32.4 pg/ml, p = 0.02) and interleukin-12 (IL-12) (0 pg/ml vs. 11.3 pg/ml, p = 0.03) at 1.5h timepoint compared to the WT mice (Fig 3A). There was no difference at the 6 h timepoint (581.7 pg/ml vs. 427.0 pg/ml, and 0 pg/ml vs. 1.8 pg/ml, respectively). We also performed qPCR quantification of the v-rel reticuloendotheliosis viral oncogene homolog A (avian) (*Rela,* aka. *p65 NFkB*) (Qin et al., 2007) and jun proto-oncogene (*Jun*) in tissues collected 6 h post induction to assess activation of the toll-like receptor 4 signalling pathway. We did not detect differences in LPS-induced upregulation of *Rela* (brain and liver) or *Jun* (liver) in FINCA mouse tissues compared to WT controls (Fig. S2). The trend in delayed IFNγ secretion was also seen in the 9-month-old mice but did not reach statistical significance (Fig. 3B). The corticosterone levels of FINCA and WT mice during LPS induction did not reveal differences in stress levels, which could have influenced the immune response (1021 ng/ml vs. 1011 ng/ml at 8 weeks of age and 1422ng/ml vs. 1350ng/ml at 9 months of age) (Fig 3). In addition, we induced experimental autoimmune encephalomyelitis (EAE) with myelin oligodendrocyte glycoprotein (MOG) to study the effect of chronic autoimmunity reaction in the context of the CNS in FINCA mice but did not detect any differences in clinical score development compared to WT litter mates (Fig. S3). Altogether, these results suggest a defect in the early stages of the LPS-induced innate immune response in FINCA mice.

**Figure 3.**
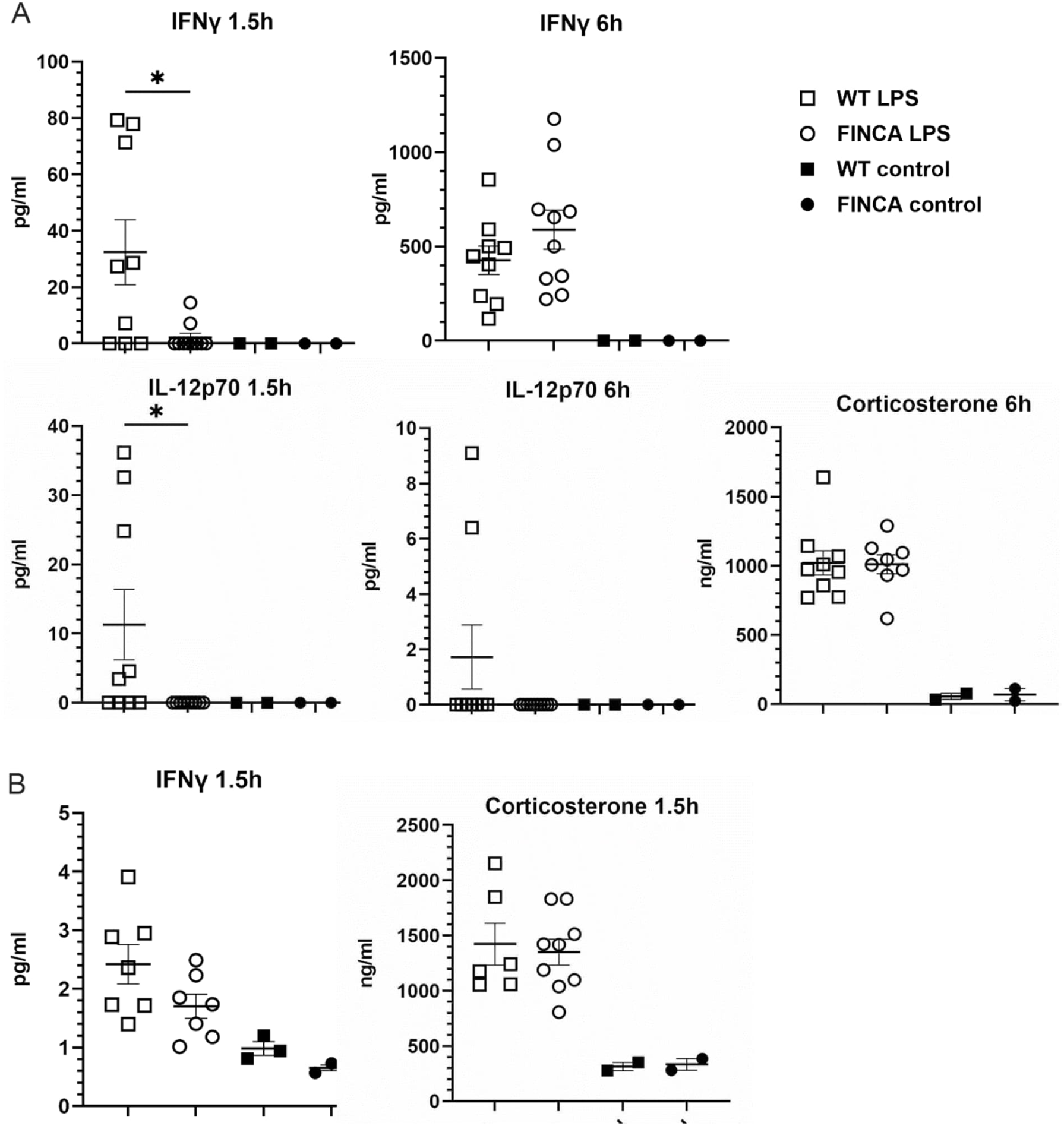
Cytokine analysis revealed a decrease in the initial IFNγ and Il-12 serum levels in FINCA mice after LPS induction. A. The cytokine response to LPS was measured 1.5h and 6h post-induction in WT (N = 9) and FINCA (N = 10) mice at 8 weeks of age. There was no difference in corticosterone levels 6 h after LPS induction in FINCA mice (N = 8) compared to WT littermates (N = 9). B. Cytokine response to LPS in WT (N = 7) and FINCA (N = 7) mice at 10.5 months of age indicated a trend towards decreased serum IFNγ 1.5h after LPS injection. No difference was detected in corticosterone levels. In terms of the statistical tests, the Mann–Whitney test was used for IFNγ, IL12p70 and corticosterone. Individual datapoints, mean and SEM are shown. * p < 0.05, ** p < 0.01, *** p < 0.001.

We further examined the FINCA mouse splenic T cells and their activation *in vitro* using flow cytometry. Representative images of the grating used for uninduced and phorbol 12-myristate 13-acetate (PMA) and ionomycin-induced splenocytes are shown in Supplementary Figures S4, S5 and S6. The analysis revealed that the FINCA mice had fewer CD25+ regulatory T cells compared to the WT mice (10.2% vs. 11.3%, p = 0.009) (Fig 4A). Cytokine analysis after PMA/ionomycin stimulation of splenocytes revealed lower amount of IFNγ-secreting (4.5% vs. 3.8%, p = 0.04) and an increased amount of interleukin 2 (IL-2)-secreting CD4+ T cells (22.0% vs. 25.2%, p = 0.02) (Fig 4B). Interestingly, uninduced FINCA splenocytes treated with only the transport inhibitor showed significantly lower amount of tumour necrosis factor (TNFα)-producing cells in both CD4+ (7.7% vs. 4.8%, p = 0.005) and CD8+ (1.72% vs. 0.87%, p = 0.0006) populations (Fig. 4C). Also, when comparing geometric means to evaluate the amount of cytokine produced, TNFα production was lower in TNFα-producing FINCA CD4+ (7536 vs. 6455, p = 0.0009) and CD8+ (9830 vs. 9059, p = 0.0166) T cells before stimulation but was statistically higher after stimulation in FINCA CD4+ cells compared to WT cells (9397 vs. 10196, p = 0.035) (Fig. 4D). After induction, a larger proportion of activated CD44+ CD8 T cells secreted TNFα in FINCA compared to WT splenocytes (60.5% vs. 51.6%, p = 0.03) (Fig 4C). Altogether, these results indicate that FINCA mice have altered regulation of T cell activation in splenocytes that leads to a decrease in IFNγ secretion and an increase in TNFα and IL-2 cytokine production.

**Figure 4.**
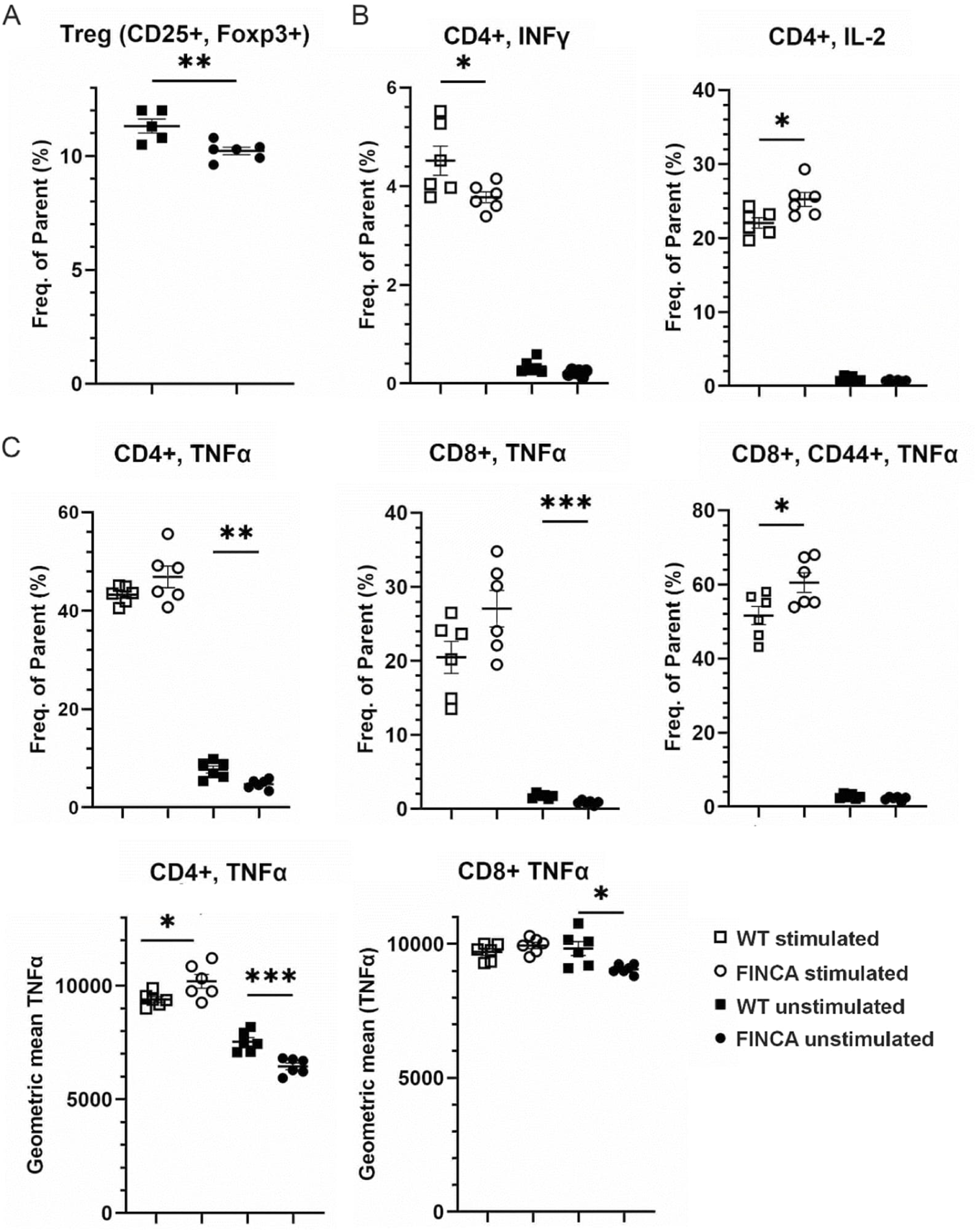
Flow cytometric analysis of stained WT and FINCA mouse splenocytes. A. FINCA mice had fewer CD25+, CD4+ regulatory T cells (Treg) compared to WT mice. B. FINCA mice showed less IFNγ-producing and more IL-2-producing CD4+ T cells after PMA/ionomycin induction. C. FINCA mice had less TNFα-producing CD8+ and CD4+ cells before PMA/ionomycin induction. After induction, FINCA splenocytes had more activated TNFα-producing CD8+, CD44+ cells. An analysis of geometric means indicated decreased TNFα production in CD4+ and CD8+ cells, as well as increased production in CD4+ cells after induction. Statistical analysis was conducted using an unpaired Student’s t-test. The mean of cells analysed from each mouse, mean of group and SEM of group are shown. * p < 0.05, ** p < 0.01, *** p < 0.001.

### FINCA male mice are less susceptible to bleomycin-induced pulmonary fibrosis

Pulmonary fibrosis, including unique granuloma-like lesions, is one of the main and often fatal manifestation of FINCA disease (Badura-Stronka et al., 2022; Rapp et al., 2021; Sczakiel et al., 2023; Tallgren et al., 2023; Uusimaa et al., 2018). Despite the significant decrease in NHLRC2 in the FINCA mouse lungs, we have not observed any abnormalities in the tissue histology or signs of granuloma-like lesions and fibrosis (Hiltunen et al., 2020). Bleomycin induction, the golden standard of modelling lung fibrosis in mice, leads to bronchiolocentric accentuated fibrotic changes, acute interstitial and intra-alveolar inflammation, macrophage activation and up-regulation of TNFα, granulocyte-macrophage colony-stimulating factor and some interleukins (Della Latta et al., 2015). Following the acute inflammatory event, other cytokines such as transforming growth factor beta and connective tissue growth factor, are up-regulated during the repair and fibrotic stage (Della Latta et al., 2015). Due to severe lung fibrosis in some FINCA patients and altered inflammatory responses in mice we hypothesised that FINCA mice would be more susceptible to bleomycin-induced fibrosis. We performed bleomycin induction on five WT males, seven FINCA males, nine WT females and seven FINCA females at 8 weeks of age. The area of fibrotic lesions and the collagen content of the lesions were quantified two weeks after bleomycin induction from Masson’s trichrome–stained lung sections. The FINCA male mice showed a smaller relative fibrotic lesion area post-induction compared to their WT counterparts (0.21 vs. 0.38, p = 0.03) (Fig. 5A). Similar trend was observed in the female mice but this effect was not statistically significant (0.17 vs. 0.3, p = 0.2). No difference was observed in the amount of collagen in the lesion sites (FINCA and WT 0.073 vs. 0.07 in males and 0.074 vs. 0.07 in females) (Fig. 5B). Based on the findings from these time points, we can conclude that FINCA malemice are less susceptible to bleomycin-induced pulmonary fibrosis.

**Figure 5.**
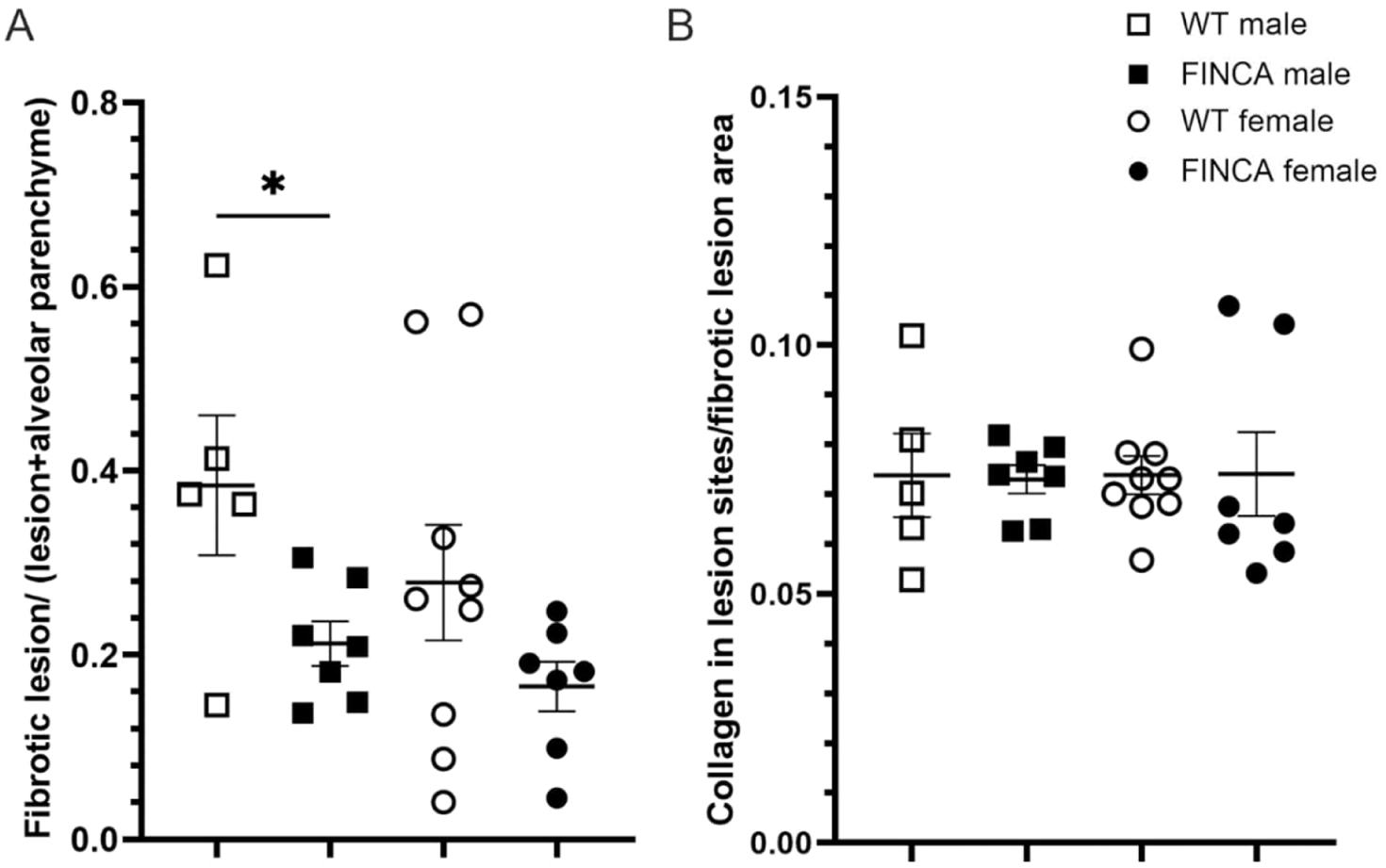
Pulmonary fibrosis in FINCA mice compared to WT littermates 14 days post bleomycin-induction. The area of fibrotic lesions, collagen in fibrotic lesion, and alveolar parenchyma were analysed using Visiopharm. A. The fibrotic lesion area was decreased in FINCA male mice (N = 7) compared to WT male mice (N = 5), and a similar trend was seen in FINCA females (N =7) compared to WT females (N = 9). B. There was no difference in the amount of collagen within the fibrotic lesions due to the genotype. Statistical analysis was done using an unpaired Student’s t-test. Individual datapoints, mean and SEM are shown. * p < 0.05.

### Proximity-labelling mass spectrometry reveals changes in putative protein– protein interactions of NHLRC2 due to the FINCA variant in HEK cells

We utilised the proximity-dependent biotin identification (BioID) (D. I. Kim & Roux, 2016) and mass spectrometry in human embryonic kidney Flp-In 293 T-REx cells to identify interacting partners of human NHLRC2 with the p.Asp148Tyr variant and compared them with the protein–protein interactions (PPIs) of wild type human NHLRC2 utilizing the constructs and cell lines established earlier (Paakkola et al., 2018). We identified 109 proteins with reduced interaction and 39 with increased interactions (p < 0.05), with the p.Asp148Tyr variant containing protein compared to WT NHLRC2 (Supp. Data 2). The most enriched pathways in the proteins with reduced interaction were ‘golgi-to-ER retrograde transport’ and ‘signaling by Rho GTPases’ in REACTOME and ‘bacterial invasion of epithelial cells’, ‘fatty acid biosynthesis’ and ‘salmonella infection’ in KEGG (Supp. Data 2). The most enriched pathways in proteins with increased interaction with the FINCA variant protein were folding by CCT/TriC in REACTOME and ‘antigen processing and presentation’, the ‘oestrogen signalling pathway’ and ‘lipid and atherosclerosis’ in KEGG (Supp. Data 2). Next, we compared these 148 proteins to the DEGs identified in FINCA mESCs to identify common pathways. We discovered 32 hits common to both datasets (Fig. 6A). Pathway enrichment analysis indicated an attenuation phase, an HSP90 chaperone cycle for steroid hormone receptors (SHR) in the presence of ligand, an HSF1-dependent transactivation, and a RHO GTPase cycle to be affected according to REACTOME and antigen processing and presentation as well as lipid and atherosclerosis according to KEGG (Suppl. Data 2). Out of the 32 common hits, 19 had reduced interaction in BioID and increased expression in FINCA mutant mESCs suggesting the possible activation of a compensatory mechanism in the mESCs. REACTOME and GO term enrichment analysis of these 19 proteins indicated RHO GTPase cycle (DST, DSP, ESYT1, DIAPH1, SLK, AKAP12 and DYNC1H1) as a compensatory pathway (Fig. 6B, Suppl. Data 2).

**Figure 6.**
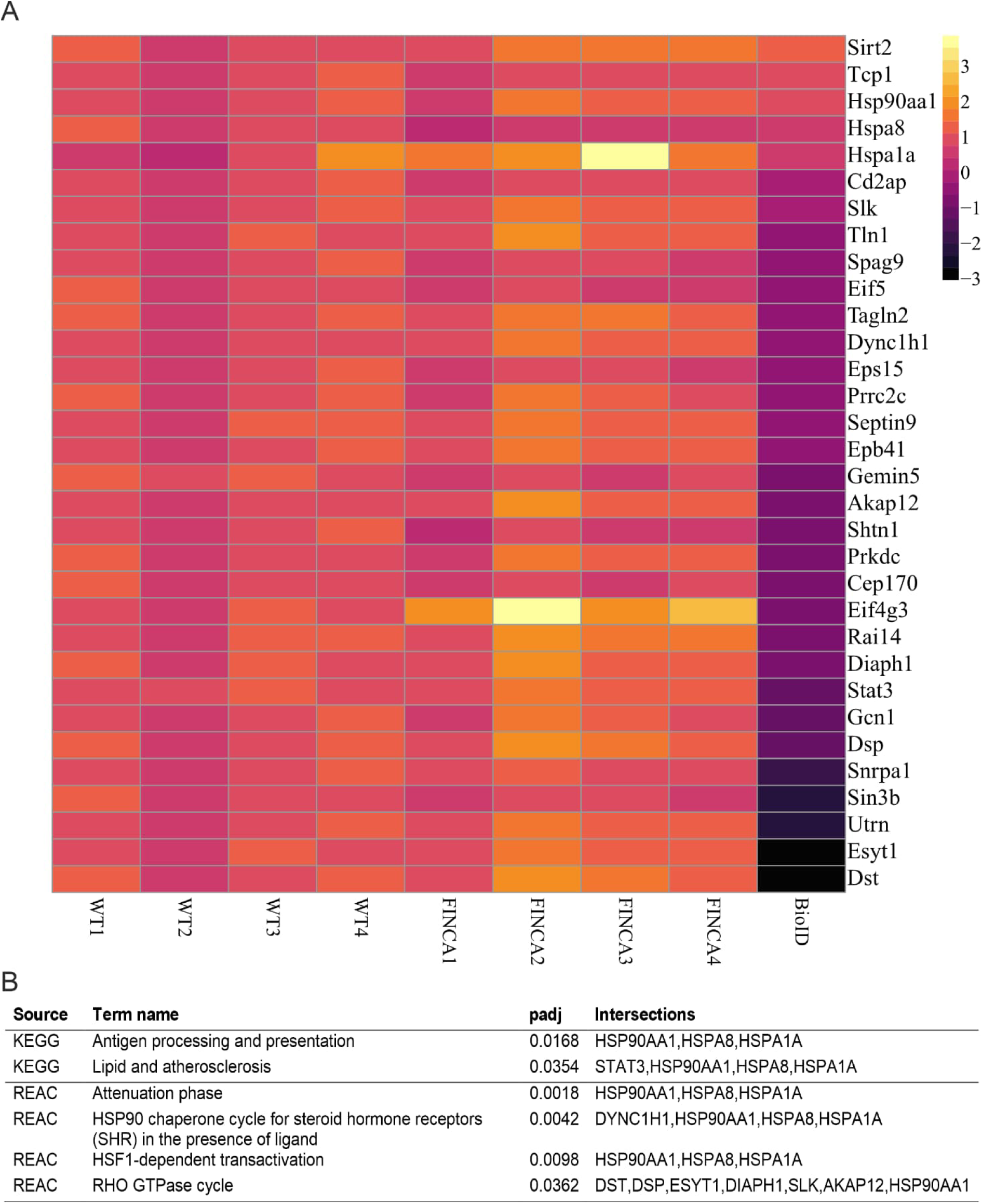
A. Heatmap of common hits when comparing the FINCA variant and WT NHLRC2 protein in mESCs with RNA sequencing and human FINCA variant and WT NHLRC2 protein in BioID. B. Pathway analysis results.

## Discussion

Knowledge of the molecular pathways affected by the disease-causing gene and the natural course of disease progression are instrumental in the development of novel therapeutics. In the current study, we elucidated molecular pathways affected by the FINCA disease-causing p.Asp148Tyr variant in NHLRC2 using both mouse and human-derived cell culture models. In addition, we performed neurobehavioral and immunological phenotyping of the FINCA mouse model to better understand the course of FINCA disease progression. We discovered ADHD-like behaviour and altered response to LPS challenge in the FINCA mouse model. At the cellular level, we report significant transcriptional remodeling associated particularly with the p.Asp148Tyr variant, which we associated with changes in the NHLRC2 interactome.

We first performed and validated RNA sequencing to compare the *Nhlrc2* KO or FINCA variant p.Asp148Tyr-expressing mESCs to WT cells. The complete loss of the gene was fairly well tolerated by mESCs and considerably larger amount of DEGs were observed in the FINCA variant cells. This is in line with the finding that NHLRC2 is not essential for mESCs (Hiltunen et al., 2022). In addition, it may indicate that, rather than pure loss of function there may be an additional gain of function or cytotoxic effects due to the missense variant requiring compensatory transcriptional changes to occur. Among others, it was verified that the expression of *Nckap1l* was decreased in *Nhlrc2* KO cells. NCKAP1L is a hematopoietic-specific WAVE complex component that regulates F-actin polymerisation downstream of RAC (Chan et al., 2013). Both NHLRC2 and NCKAP1L have been associated with phagocytosis in macrophages (RAW 264.7 cells) (Haney et al., 2018). NCKAP1L has been shown to be involved in lymphocyte development, activation, proliferation and homeostasis, erythrocyte membrane stability, as well as phagocytosis and migration by neutrophils and macrophages (Chan et al., 2013; Park et al., 2008). Recently, according to a conference abstract, NHLRC2 has been associated with human erythroid development (Myers et al., 2023), which would explain the haemolytic macrocytic anaemia described in FINCA patients (Tallgren et al., 2023). In human T cells, NCKAP1L is required for the proper mechanistic target of rapamycin complex 2 (mTORC2)-dependent AKT phosphorylation, cell proliferation and cytokine secretion, including IL-2 and TNFα (Cook et al., 2020) secretion, which we found to be affected in the FINCA mouse model splenocytes (Fig. 3 and 4). The expression of *Nckap1l* was significantly decreased only in *Nhlrc2* KO mESCs suggesting that the association may be stronger related to the physiological function of NHLRC2 rather than pathological consequence of p. Asp148Tyr variant .

It was verified that *Slc27a2* expression was decreased in both the *Nhlrc2* KO and FINCA variant mESCs. SLC27A2 (aka FATP2, VLCAS) mediates the import of long-chain fatty acids into the cell and is a peroxisomal very long-chain acyl-CoA synthetase (Hirsch et al., 1998; Uchiyama et al., 1996). SLC27A2 regulates non-small-cell lung cancer (Chen et al., 2023) and kidney fibrosis (Chen et al., 2020) through its effect on lipid metabolism. A high level of NHLRC2 has been associated with shortened survival in lung adenocarcinoma, a type of non-small-cell lung cancer (Kreus et al., 2023). In the neurological context, it has been suggested that impaired SLC27A2 activity leads to the accumulation of saturated very-long-chain fatty acids in a demyelinating disorder X-adrenoleukodystrophy (MIM: 300100) (Smith et al., 2000). At the pathway level, neurodevelopmental and collagen synthesis pathways were affected in FINCA-variant mESCs, which nicely correlates with the neurodevelopmental disorder and fibrosis observed in patients (Tallgren et al., 2023; Uusimaa et al., 2018).

FINCA patients have been reported to manifest with neurodevelopmental disorder from early infancy, intellectual disability, severely retarded speech, relatively preserved speech comprehension, behaviour problems (e.g. aggression out-bursts, attention deficit and irritability), epileptic seizures, muscle weakness and movement disorder (Badura-Stronka et al., 2022; Rapp et al., 2021; Sczakiel et al., 2023; Tallgren et al., 2023; Uusimaa et al., 2018). Additionally, neural tube-related developmental malformations called development duplications (DD, OMIA 002103-9913) are associated with the p.(Val311Ala) variant in Angus cattle underscoring the importance of NHLRC2 in central nervous system development (Polkoff, 2017). Here, we conducted a comprehensive behavioural phenotyping that revealed mild hyperactivity in FINCA mice compared to WT littermates at 13 weeks of age, which had abated by 9 months of age. These findings suggest an ADHD-like neurodevelopmental defect in FINCA mice, partly recapitulating the patient phenotype. However, the phenotype was considerably milder compared to that seen in FINCA patients, indicating that there is a species and/or environmental aspect influencing the severity of this disorder. Furthermore, future studies will clarify whether there are less severe variants of NHLRC2 behind milder forms of neurodevelopmental defects in humans. For example, in one recent publication, *NHLRC2* was found to correlate with a behavioural subtype of autism spectrum disorder (Q. Guo et al., 2023).

Recurrent infections are a common symptom of FINCA in early childhood and lead to worsening of the disease state (Badura-Stronka et al., 2022; Rapp et al., 2021; Sczakiel et al., 2023; Tallgren et al., 2023; Uusimaa et al., 2018). We assessed the immune response of FINCA mice *in vivo* by systemic LPS injection and serum cytokine analysis, as well as FINCA mouse splenocytes *in vitro* by intracellular cytokine staining and T cell analysis using flow cytometry (Fig. 4). We found IFNγ response to be delayed and IL-12 response to be absent in FINCA mouse serum after LPS induction. FINCA splenocytes indicated the dysregulation of IFNγ, IL-2, and TNFα production. Our findings also revealed a possible increase in TNFα in FINCA mouse T cells already at baseline, which could lead to a disturbed immune response. IFNγ, IL-2, and IL-12 mediate the proinflammatory Th1 immune responses, where IFNγ is the main Th1 cytokine. In thymocytes, NHLRC2 has previously been reported to be a target gene for nuclear zinc-finger protein Zfat, a transcription factor suggested to be important for T cell survival and development in mice (Ishikura et al., 2015, 2016). Immune dysfunction is commonly associated with several neurological and mental disorders, including autism spectrum disorder (Ashwood et al., 2011). In particular, IFNγ has been found to regulate neuronal survival and connectivity and to affect social behaviour (Filiano et al., 2016). Our findings indicate that the FINCA mouse model has altered innate immune response and T cell−mediated cytokine secretion in connection with mild neurodevelopmental disorder.

FINCA patients often manifest with pulmonary fibrosis (Badura-Stronka et al., 2022; Rapp et al., 2021; Sczakiel et al., 2023; Tallgren et al., 2023; Uusimaa et al., 2018). IFNγ has been shown to ameliorate bleomycin-induced lung fibrosis in mice (Giri et al., 1986), and thus, we expected FINCA mice to be more susceptible to bleomycin-induced lung fibrosis. However, FINCA male mice indicated a decreased formation of fibrotic lesions after bleomycin administration compared to WT male mice. More information on the temporal changes in both cytokine response and fibrogenesis is needed to fully understand the role of NHLRC2 in pulmonary fibrosis.

Finally, to identify common conserved pathways between mice and humans, we compared significant DEGs identified in FINCA variant mESCs with protein−protein interaction partners affected by the FINCA variant in human NHLRC2 that were identified by BioID. Decreased interaction was seen in Golgi-to-ER retrograde transport and Rho GTPase signalling-associated proteins. We have previously shown that NHLRC2 affects vesicle transport in human fibroblasts (Paakkola et al., 2018). Additionally, the retromer component sorting nexin-6 (SNX6) was identified as a putative interaction partner of NHLRC2 in HEK 293 cells (Paakkola et al., 2018) and (SNX6) was later shown to decrease in mouse NPCs due to the FINCA variant (Hiltunen et al., 2020). The identification of Rho GTPase signalling pathway is consistent with a previous report in which the loss of NHLRC2 was found to inhibit phagocytosis, possibly through Rho/Rac1 signalling (Haney et al., 2018). Rho GTPase signalling has emerged as an important regulatory pathway in a number of conditions relevant to FINCA disease, including neurodevelopmental disorders (Banka et al., 2022; D. Guo et al., 2020) and immune syndromes (El Masri & Delon, 2021).

In conclusion, our data show that FINCA mice have symptoms similar to human patients, including neurodevelopmental and immunological disorders, confirming their feasibility as a model organism for studying the pathological processes of NHLRC2. Our results reveal the first hints of cytokine cascades affected by deficient NHLRC2 in mice. Future studies will elucidate whether targeting specific immune responses in patients can alleviate their disease course.

## Materials and methods

### Study subjects and materials

#### Mice (Mus musculus)

The FINCA mouse model used in this study was a hetero compound progeny of C57BL/6NCrl-*Nhlrc2*^tm1a(KOMP)Wtsi^ (EMMA ID: EM:10219) (Skarnes et al., 2011) (*Nhlrc2* KO) and C57BL/6NCrl-*Nhlrc2*^em1Rthl^ (Hiltunen et al., 2020) mice. Both mouse lines have been archived to the European Mouse Mutant Archive (https://www.infrafrontier.eu/emma/). WT littermates were used as controls. Animal care and experimental procedures were conducted in adherence to European Union regulations and guidelines and the Finnish legislation on housing laboratory animals (DIRECTIVE 2010/63/EU, Act 497/2013 and Decree 564/2013). Animal experiments conducted in Finland were approved by the Project Authorisation Board of Finland (ESAVI/33827/2019, ESAVI/14247/2019). Bleomycin induction was performed at the Oulu University Animal Centre. Behavioural studies and LPS induction on 9-month-old mice were conducted at the Lab Animal Centre of University of Eastern Finland. All of the conditions of LPS responses at 8 weeks of age and EAE studies, performed by Biomedcode (Vari Greece), conformed to Presidential Decree No 56/2013 (Governmental Gazette No A’ 106), which is applicable in Greece and is the implementation of the EEC Directive 2010/63/EEC. They were approved by the Directorate of Agricultural and Veterinary Policy (DAVP) of the Attica Region (approval license protocols nos. 159982/26-02-2021 and 159971/26-02-21 respectively).

#### Mouse embryonic stem cell culture

Mouse mESC lines were established from four WT, four *Nhlrc2* KO (C57BL/6NCrl, homozygous for *Nhlrc2*^tm1a(KOMP)Wtsi^ allele), and four FINCA (C57BL/6NCrl, homozygous for *Nhlrc2*^em1Rhtl^ allele) mouse blastocysts at the Biocenter Oulu Transgenic and Tissue Phenotyping Core Facility at the University of Oulu, Finland, as previously described (Hiltunen et al., 2022; Nichols et al., 2009). The cells were maintained on gelatinised plates in complete basal medium (Millipore, Billerica, MA, USA) supplemented with a 2i Supplement Kit (Millipore, Billerica, MA, USA) and 100 u/ml penicillin/streptomycin (Sigma-Aldrich, St. Louis, MO, USA). The cells were grown in a cell culture incubator at 37°C and 5% CO_2_. The medium was changed daily, and the cells were passaged using Accutase (Millipore, Billerica, MA, USA).

#### Human embryonic kidney cell culture

Flp-In^tm^293 T-Rex (HEK 293T, Thermo Fisher Scientific, Waltham, MA, USA) cells were maintained in high-glucose DMEM (Corning, Manassas, VA, USA) supplemented with 10% FBS in a cell culture incubator at 37°C and 5% CO_2_.

## Methods

### RNA isolation from mouse embryonic stem cells and tissues

RNA isolation from the mESCs was performed using an RNeasy Mini Plus Kit (Qiagen, Hilden, Germany) and an RNase-Free Dnase Set (Qiagen, Hilden, Germany). RNA isolation from mouse liver samples was carried out using an RNeasy Plus Mini Kit (Qiagen, Hilden, Germany) and from brain samples using an RNeasy Lipid Tissue Mini Kit (Qiagen, Hilden, Germany) according to the manufacturer’s instructions.

### RNA sequencing

The quality control and RNA sequencing of four *Nhlrc2* KO, four FINCA variant, and four WT mESCs were performed at the Biocenter Oulu Sequencing Center, Oulu, Finland. Precise concentrations were measured with Qubit Assay, and Bioanalyzer Assay were used to determine RNA integrity. The library was prepared using RNA seq, TruSeq^®^ Stranded Total RNA Kit (Illumina, San Diego, CA, USA). The sequencing was performed in pair-end mode with the NextSeq 550 High Output v2.5 Kit (150 cycles). Each sample was sequenced across three lanes.

### RNA data analysis

Before starting the analysis, the multiple lanes were merged to generate a “master” fastq file per sample.

The DEGs were analysed using Chipster (Kallio et al., 2011). In short, quality control of the fastq files was performed using FastQC (Wingett & Andrews, 2018). Alignment was performed using HISAT2 (D. Kim et al., 2015) for paired end reads with reference genome *Mus_musculus*.GRCm38.95. Alignment quality control was performed with RseQC (Wang et al., 2012). HTSeq was used for quantification (Anders et al., 2015), and DEGs were detected using DESeq2 (Love et al., 2014) where the p-values attained by the Wald test are corrected for multiple testing using the Benjamini and Hochberg method. Adjusted p-value less than 0.05 were considered statistically significant.

For alternative splicing analysis it is crucial that all the reads have same read length (Gondane & Itkonen, 2023). FastQC analysis revealed that the “master” fastq files had variable read length and thus, trimmomatic was used to trim the fastq files so that each read was 76 bp long (Bolger et al., 2014). rMATS v2.7 (Shen et al., 2014) was used to call the differentially spliced sites using the *Mus-musculus*.FRCm39 . A bar plot depicting the count of splicing events was made using the ggplot2 package in RStudio to show the events that were significant (p < 0.05).

### Quantitative PCR

QPCR was used to verify selected RNA sequencing hits. Validation was performed on RNA samples collected from three FINCA variant, three KO and three WT mESC clones. The cells were cultured, and RNA isolation was performed separately from the RNA sequencing samples using the same protocol as when preparing samples for sequencing. A QuantiTect Reverse Transcription Kit (Qiagen, Hilden, Germany) was used for complementary cDNA synthesis. Primers (Suppl. Table 1) were designed using NCBI Primer-BLAST (Ye et al., 2012), and Tm was set at 60°C. QPCR was performed according to the manufacturer’s instructions (IQTM SYBR Green Supermix, Bio-Rad, Hercules, CA, USA) with a CFX Connect^TM^ Real-Time System (Bio-Rad, Hercules, CA, USA). Expression analysis was performed using the Bio-Rad CFX Maestro program.

### Mouse behavioural tests

Behavioural tests were conducted on 16 WT and 14 FINCA female mice at 2.5 and 9 months of age at the University of Eastern Finland in the order described below.

#### Exploratory activity

TruScan^®^ (Coulbourn Instruments, Allentown, PA, USA) automated activity monitoring, based on infrared photobeam detection was used to assess spontaneous locomotion and exploratory activity. The system comprises a white-walled observation cage (26 × 26 × 39 cm) and two rings of photobeam detectors to monitor horizontal and upright positions. A free 10-min exploration was recorded. The analysed parameters included ambulatory distance (gross horizontal locomotion), stereotypy time (time engaged in movements that repeatedly break adjacent beams three times), time in the cage centre and vertical time (rearing).

#### Nest-building

The nest-building test was used to evaluate each mouse’s ability to carry out goal-oriented motor sequences. The mouse was placed in an individual cage, an all enrichment material was removed and replaced with a compact cotton pad (Nestlet). The mouse was freely left to build a nest overnight. The outcome was photographed and scored by blinded raters according to the original description (Deacon, 2006) on a scale of 0–4.

#### Elevated plus maze

The elevated plus maze test is a widely used measure of anxiety. A maze comprising four arms (25 × 5 cm) radiating from a central platform (5 × 5 cm) was placed 50 cm above the floor. Two of the arms were open and two were closed with a 16 cm high wall. The maze was made of white plastic for contrast in the video image. In each test, a mouse was placed on the central platform with its nose pointed to a closed arm, and it was video-recorded for 5 min. The number of transitions between the arms and the time spent in the open and closed arms were calculated. The percentage of total time spent in open arms was analysed.

#### The Marble burying task

is another widely used test for anxiety and especially object neophobia. In each test, the mouse continued to stay in an individual cage. Nine marbles (1 cm diameter) were placed on one-half of the cage in a 3 × 3 array, and approximately 0.8L of clean bedding material was added to the cage to help bury the marbles. The marbles were left overnight, and next morning, the number of visible marbles was counted. A high number of visible marbles (unless all marbles had been left untouched) indicate high anxiety.

#### Grid hanging

This test was used to determine static muscle force. Each mouse was placed on a 20 × 25 cm wire grid (grid unit 1 × 1 cm) that was carefully placed upside down as the lid of a 24 × 35.5 × 24 cm transparent plastic cage. The time to fall off until a cut-off time of 300 s was measured with a stopwatch. The test was repeated three times with a 10-min interval between the trials and the best result was recorded. After the test, the mouse was returned to a group cage.

#### Social approach

A social approach test was performed to analyse possible deviations in the social behaviour of the mice. The test was carried out in a metal cage (43 × 25 × 15 cm) with bedding and two cylindrical cages (height 9 cm, diameter 10 cm, with filled 0.5 L plastic bottles on the top to add weight) on opposite ends of the cage. Two opaque plastic walls separated the lateral compartment from the cage, with a neutral zone in the middle. A mouse-sized opening in the partition allowed free access between compartments. After a 10-min habituation period, a stranger mouse (one mouse from the study group) was placed in one of two cages. The actual test took 10 min and was video-recorded. The number of nose contacts with the cages and the time sniffing were calculated offline from the recording by a grader blinded to the mouse genotype.

#### Rotarod

The Rotarod test was performed to assess motor coordination and balance. Mice were tested using an automated Rotarod device (Hugo Basile Rota-Rod for mice, Italy). On the first day, the mice were left to adapt to staying on the rod at the lowest speed (5 rpm) for 2 min. The next day, the test started at 5 rpm, and the speed was increased in 30 s steps until 40 rpm. The cut-off was set to 6 min. If the mouse fell or rotated with the rod for three rounds without any correcting steps, the test was stopped and the time was recorded.

#### A hot plate

was used to assess the threshold for thermal pain. Each mouse was placed on a hotplate (IITC Life Science, Woodland Hills, CA, USA) and the time to lick a paw or jump was recorded until a cutoff time of 60 s. The mouse was tested three times a day, with 1 h between trials. On day 1, the surface temperature was 47°C, and on day 2, it was 51°C.

#### Novelty-suppressed feeding

Each mouse was placed overnight in an individual cage with new bedding and the amount of food pellet was limited to one-third of the daily consumption. The next day, the mouse was placed in a new arena (66 × 48 × 28 cm) with a single standard pellet at the far end of the mouse. The time to sniff and bite the pellet was taken with a stopwatch.

#### Morris swim task

(water maze). The Morris swim task was used to measure spatial learning and memory. The apparatus was a white plastic wading pool with a diameter of 120 cm filled with water and located in a room containing distal visual cues on the walls. A transparent glass platform (10 × 10 cm) was placed 1.0 cm below the water surface. The temperature of the water was maintained at 20 ± 0.5°C throughout the experiment, and a short recovery period in a warm cage was allowed between the trials. First, each mouse was pretrained for two days to find and climb onto the submerged platform, which was placed between the southwest and northwest quadrants, by guiding it through an alley with high walls. During the spatial learning phase (days 1–4), the animals were given five trials/day with 10 min of rest between the trials. On day 5, the platform was removed from the pool for the first and last trials to determine search bias. The platform location was kept constant, but the starting position varied between four constant locations at the pool rim (north, west, south and east), with all mice starting from the same position in any single trial. If a mouse failed to find the escape platform within 60 s, it was placed on the platform for 10 s by the experimenter. The task was video-recorded with a ceiling camera. The escape latency and swimming speed were calculated for each trial and then averaged across all 5 days of task acquisition. The search bias during the probe trial was measured by calculating the time the mice spent within 30 cm from the centre of the former platform position.

#### Novel object recognition

This test was used to evaluate recognition memory. At the age of 2.5 months, half of the mice were given object A (a 50 mL test tube with a yellow paper strip inside) and the other half were given object B (the piston of a standard plastic 20 mL syringe) to familiarise themselves with the object in their home cages (with all enrichment removed) for 20 min. The next day, the mice were given 10 min to explore the same objects. Touching the object with the nose or the nose pointing to the object closer than 1 cm was counted as object exploration. Mice that did not spend 20 s in object exploration were excluded from the test. The test phase started 60 min later. The familiar object (A or B) was paired with a new object (B or A), with the position balanced within the groups of animals. Exploration of the new versus the familiar objects was video recorded for 5 min. At 9 months of age, the test was repeated with two new objects, a metal doorknob (C) and a green square Duplo block (D). Novelty preference (%) was calculated as follows: (time at the new object - time at the familiar object) * 100 / total exploration time.

#### Forced swimming

This test is widely used to assess behavioural despair. Two 3.0 L glass containers were placed on a table with a wall to block the view between them. The containers were filled with 2.0 L of water (T 22°C). Two mice were tested at the same time. The mice were gently placed on the water surface for 6 min and video-recorded. Immobility (floating) time during the last 4 min was calculated offline by a blinded rater. After the test, the mice were dried with a towel and placed under an infrared lamp to warm up.

### LPS induction and serum analysis

Eleven WT and 12 FINCA female mice were transferred to Biomedcode Greece at 6 weeks of age. At 8 weeks of age, nine WT and 10 FINCA were treated with 10 μg/mouse LPS from *Escherichia coli* (O55:B5 MD Bioproducts) administered intraperitoneally (ip). Two WT and two FINCA mice were kept untreated to serve as controls. Sera were collected 1.5 h and 6 h post-induction and subjected to cytokine and corticosterone measurements. IL-23, IL-1α, IFNγ, TNFα, MCP-1, IL-12p70, IL-1β, IL10, IL27, IL-27, IL-7A, IFNβ and GM-CSF were measured from serum samples using multiplex ELISA (LEGENDplex™ Mouse Inflammation Panel). IL-6 and CXCL1 levels were measured using Quantikine ELISA (M6000B-1 and MKC00B-1, respectively). Brain and liver tissues were collected for RNA isolation at 6h post LPS-induction into RNAprotect (Qiagen, Hilden, Germany).

For mice undergoing behavioural tests, LPS (Sigma-Aldrich, Saint Louis, MO, USA) induction was done after completion of the behavioural tests at the age of 9 months. Seven WT and eight FINCA females were treated with 18 ug/10g of LPS ip. Three mice from each genotype were treated with the vehicle alone. Sera were collected 1.5 h after LPS induction for cytokine measurements. Cytokines (Suppl. Table 2) were analysed using the U-PLEX assay (Meso Scale Discovery) according to the manufacturer’s instructions. Serum samples were diluted at a ratio of 1:4 and, 25 µl of the dilutions was added to a U-PLEX well for duplicates.

Corticosterone was measured from the serum samples of all LPS-induced mice and their controls using a competitive enzyme-linked immunosorbent assay kit (ab108821, Abcam, Cambrige, UK) according to the manufacturer’s instructions. Measurements were taken with a FLUOstar Omega microplate reader (BMG LabTech, Ortenberg, Germany).

### MOG induced EAE

Eight-week-old male mice (N = 8 WT and N = 10 FINCA) were subjected to the EAE induction protocol. More specifically, on day 0, the mice were treated with 150 ng pertussis toxin (Sigma-Aldrich; P2980) ip and immunised by subcutaneous administration of an emulsion containing 150 µg MOG 35-55 peptide (peptide sequence MEVGWYRSPFSRVVHLYRNGK, GeneCust) and 1mg mycobacterium tuberculosis (strain H37Ra, BD™ Difco™ 231141) in incomplete Freund’s adjuvant (Sigma-Aldrich; F5506). On day 2, the mice were administered a second dose of 150 ng pertussis toxin to induce the opening of the blood–brain barrier. Body weight was recorded twice weekly for the entire duration of the study. On days 7–21, post-immunisation pathology was scored daily. On days 23–31 post-immunisation pathology was scored during the weekdays. Pathology scoring was based on macroscopic observations for signs of paralysis according to Supplementary Table 3.

### Mouse splenocyte isolation and FACS analysis

Splenocytes were harvested from 6 FINCA and 6 WT female mice for the analysis of T cell populations, T cell activation, and cytokine production by flow cytometry according to the protocol described earlier (Skordos et al., 2021). In short, 1.5 million splenocytes per mouse were stained for extracellular markers and fixed and stained for intracellular markers (Suppl. Table 4). Another 1.5 million cells were stimulated using eBioscience Cell stimulation (PMA/ionomycin) and inhibitor cocktails (Suppl. Table 5) and stained for extracellular and intracellular markers. Single-stained cell controls were prepared for each antibody used. UltraComp eBead Compensation Beads (Thermo Fisher Scientific, Waltham, MA USA) were prepared for each antibody except for the fixable viability dye. An amine reactive compensation bead kit was used (Thermo Fisher Scientif, Waltham, MA USA) according to the manufacturer’s instructions. Fluorescence-assisted cell sorting was performed using LSRFortessa (BD Biosciences, Franklin Lakes, NJ, USA), auto-compensation was performed, and data analysis was conducted using FlowJo version 10.8.1.

### Bleomycin induction

Bleomycin (2 U/kg; 1,2 U/ml) was administered to the airways of 13 WT (five males and nine females) and 16 FINCA mice (seven males and seven females) at 8 weeks of age under isoflurane anaesthesia. The lungs were harvested 14 days post-induction. The left lobe was filled with fixative through tracheal canulation for histological analysis and samples from the right lobes were collected for RNA and protein analyses. Sections of the left lobe measuring 5µm were Masson’s trichrome stained and two sections, on both sides of the hilum and 435 µm apart, were chosen for analysis. Histological slides were scanned using a Hamamatsu NanoZoomer S60 slide scanner at 40× magnification. Whole slide images were analysed using Visiopharm software (Visiopharm, Hoersholm, Denmark) to quantify the proportion of fibrotic lesions in the total lung parenchyme and the collagen content of the fibrotic lesions.

### Proximity-dependent biotin identification

BioID analysis was performed as previously described (Paakkola et al., 2018). Briefly, previously generated human NHLRC2 tagged C-terminally with BirA-FLAG was utilized to generate the tagged FINCA variant using site-directed mutagenesis (Zheng et al., 2004).

Stable cells with the genomic insertion of recombinant wildtype NHLRC2 and FINCA variant were generated as Flp-In 293 T-REx HEK 293 cell pools according to the manufacturer’s instructions (Invitrogen, Carlsbad, CA, USA). Expression of the recombinant protein was induced using tetracycline (1µg/ml) and the cell were treated with 50 µM biotin for 24h before sample collection. The biotinylated proteins were isolated using Strep-Tactin beads (IBA, Gmbh) and analysed using liquid chromatography-MS (LC-MS). The samples were collected in triplicate. Interaction partners of the wildtype NHLRC2 were compared to the interaction partners of the NHLRC2 FINCA variant to identify potentially altered interactions caused by the variant.

### Mass spectrometry analysis

AnOrbitrap Elite ETD hybrid mass spectrometer (Thermo Fisher Scientific, Waltham, MA) with an EASY-nLC II system was used for the LC-MS/MS analysis. Peptide separation was performed with a precolumn (C18-packing; EASY-Column™ 2 cm × 100 μm, 5 μm, 120 Å, Thermo Fisher Scientific) followed by an analytical column (C18-packing; EASY-Column™ 10 cm × 75 μm, 3 μm, 120 Å, Thermo Fisher Scientific). A 60-minute linear gradient from 5–35% buffer B (Buffer A: 0.1% formic acid in HPLC-grade water; buffer B: 0.1% formic acid in 98% acetonitrile and 2% HPLC-grade water) was used at a stable flow rate of 300 nl/min. Data-dependent acquisition was performed using one high-resolution (60000) FTMS full scan (m/z range 300–1700), followed by the top 20 CID-MS2 scans (energy 35). The highest fill time was 200 ms for FTMS and 200 ms for the ion trap. Precursors with ion counts of over 500 were chosen for MSn, and preview mode was applied for the FTMS scan to achieve a high resolution. Dynamic exclusion was enabled with repeat count setting of 1, a repeat duration of 30, an exclusion list size of 500, an exclusion duration of 30, an exclusion mass width relative to low (ppm) of 5.000 and an exclusion mass width relative to high (ppm) of 5.000.

### Protein identification and filtering

Protein identification from MS2 spectral data was performed using the SEQUEST search engine in Proteome Discoverer™ software (version 1.4, Thermo Fisher Scientific). The raw data were analysed against the reviewed human proteins from the UniProt database (October 2018). Two missed cleavages and 15 ppm monoisotopic mass error were permitted, and the peptide FDR was set to <0.05. Precursor mass tolerance and fragment mass tolerance were set to 15 ppm and 0.05 Da, respectively. Lysine biotinylation and oxidation of methionine were set as dynamic modifications, and cysteine carbamidomethylation was set as a static modification. The identified proteins were filtered using SAINTexpress version 3.1.0 with a BFDR cutoff of 0.05. Further filtering was performed based on the CRAPome database (version 2.0). All interactions, where the prey protein was present with an occurrence of 20 % or more were discarded, unless their fold change against CRAPome average was three times higher or more. Spectral counts of the filtered interactions were compared using a t-test, and p-values were adjusted using FDR correction.

### Data-analysis

Pathway analyses and gene ontology enrichment analyses were performed using gProfiler (e111_eg58_p18_30541362) (Raudvere et al., 2019; Reimand et al., 2007). The data versions used were KEGG FTP Release 2024-01-22, BioMart classes: 2024-1-25 for REACTOME, and BioMart releases/2024-01-17 for GO term enrichment analysis.

### Statistical analysis

Statistical analysis and correction for multiple comparisons was performed on RNA sequencing and proximity-labelling mass spectrometry as described in corresponding methods sections. All other statistical analysis was performed using GraphPad Prism 10 (GraphPad Software, San Diego, CA, USA). Where the data exhibited a normal distribution, unpaired Student’s t-test was used for comparison. The Mann–Whitney U-test was used when the data did not follow a normal distribution to ascertain statistical significance. A p-value of less than 0.05 was considered statistically significant.

## Acknowledgements

The authors thank Dr. Helmut Pospiech, Pirjo Keränen, and Espen Poulsen for their expert assistance. Biocenter Finland, Biocenter Oulu Sequencing Centre, Biocenter Oulu Transgenic and Tissue Phenotyping Core Facility, Oulu Laboratory Animal Centre, Biocenter Kuopio Phenotyping Centre and Lab Animal Centre at the University of Eastern Finland are acknowledged for their technical support and service. We thank Translational cell biology core facility at Kontinkangas campus for their assistance with flow cytometry service. The authors wish to acknowledge CSC – IT Center for Science, Finland, for the computational and INFRAFRONTIER for mouse resources.

## Data availability

All relevant data can be found within the article and its supplementary information. Raw data and total read counts of the RNA sequencing experiment are available at Array Express (https://www.ebi.ac.uk/biostudies/arrayexpress) with accession E-MTAB-14174. Mass spectrometry data concerning the BioID experiments is available at the MassiVE repository (ftp://massive.ucsd.edu/v08/MSV000094956/).

